# Detection of precisely edited CRISPR/Cas9 alleles through co-introduced restriction-fragment length polymorphisms

**DOI:** 10.1101/2021.04.20.440537

**Authors:** Chon-Hwa Tsai-Morris, Sydney Hertafeld, Yvonne Rosario, James Iben, Eric Chang, Ling Yi, Steven L. Coon, Stephen G. Kaler, Ryan Dale, Benjamin Feldman

**Affiliations:** NICHD Zebrafish Core, *Eunice Kennedy Shriver* National Institute of Child Health and Human Development (NICHD), NIH, Bethesda, MD 20892, USA; Bioinformatics and Scientific Programming Core, NICHD, NIH, Bethesda, MD 20892, USA; NICHD Molecular Genomics Core, NICHD, NIH, Bethesda, MD 20892, USA; Section on Translational Neuroscience, NICHD, NIH, Bethesda, MD 20892, USA; Center for Gene Therapy; Abigail Wexner Research Institute, Nationwide Children’s Hospital, The Ohio State University College of Medicine; Columbus, OH 43205

## Abstract

CRISPR/Cas9 is a powerful tool for producing genomic <u>in</u>sertions and <u>del</u>etions (indels) to interrogate gene function. Modified CRISPR/Cas9 protocols can produce targeted genetic changes that are more precise than indels, but founder recovery is less efficient. Focusing on producing missense mutations in zebrafish using <u>s</u>ingle-<u>s</u>tranded <u>o</u>ligo <u>d</u>eoxy<u>n</u>ucleotide (ssODN) donor templates, we pioneered a strategy of adding synonymous changes to create novel <u>r</u>estriction-<u>e</u>nzyme (RE) sites, allowing detection of rare precise edits in a modified fluorescent-PCR fragment assay. We have named this process TIARS (<u>t</u>est for <u>i</u>ncorporation of <u>a</u>dded <u>r</u>ecognition <u>s</u>ites). Aided by TIARS, we induced two distinct amino-acid substitutions (T979I and P1387S) in the *atp7a* gene among somatic tissues of CRISPR-Cas9-treated F_0_ zebrafish. One of these F_0s_ transmitted the allele to *atp7a^T979I/+^* F_1_ progeny, and trans-heterozygosity of this allele against a null *atp7a* allele causes hypopigmentation, consistent with more severe pigment deficits in zebrafish or humans carrying only null mutations in *atp7a/ATP7A*. Design of ssODNs with novel RE recognition sites is labor-intensive, so we developed an *in silico* tool, TIARS Designer, and performed bioinformatic validation indicating that TIARS should be generalizable to other genes and experimental systems that employ donor template DNA.

## INTRODUCTION

<u>C</u>lustered <u>r</u>egularly <u>i</u>nterspaced <u>s</u>hort <u>p</u>alindromic <u>r</u>epeats/Cas9 endonuclease (CRISPR/Cas9)-based mutagenesis is a popular and successful tool for generating knock-out models with disruptive <u>in</u>sertions and <u>del</u>etions (indels) (1). Two essential components for CRISPR/Cas9 mutagenesis are 1) the Cas9 protein itself and 2) a gene target-specific <u>CR</u>ISPR RNA (crRNA) that is fused or hybridized to a Cas9-specific <u>tr</u>ans-<u>ac</u>tivating RNA (tracRNA). The crRNA/tracRNA hybrid is also called a <u>s</u>ynthetic <u>g</u>uide RNA (sgRNA), or simply gRNA. These two components are delivered to cells or embryos to effect mutagenesis. Targeting and mutagenesis is restricted to genomic sequences immediately adjacent to a so-called <u>p</u>roto-spacer <u>a</u>djacent <u>m</u>otif (PAM), which for the popular Cas9 from *Streptococcus pyogenes*, is the sequence NGG. When the gRNA/Cas9 ribonuclear protein complex arrives at its target, it is able to create a <u>d</u>ouble-<u>s</u>trand <u>b</u>reak (DSB) upstream of the PAM site. Such DSBs may be repaired perfectly via <u>n</u>on-<u>h</u>omologous <u>e</u>nd <u>j</u>oining (NHEJ), but erroneous NHEJ also frequently occurs, introducing indels, and indels that also shift the reading frame typically cause complete or partial loss of gene function.

It has more recently been discovered that, beyond random indels, one can generate precise genome edits near a target by adding a DNA repair template as a third component of the CRISPR/Cas9 mix, leading to the incorporation or copying of repair template sequence containing the edit into the genome with no other changes (2,3). Homology-based DNA repair templates contain sequence identical to the target sequence except for desired substitutions, insertions or deletions. In the zebrafish model, which is our focus here, success has been reported for both <u>s</u>ingle-<u>s</u>tranded <u>o</u>ligo <u>DN</u>A (ssODN) and <u>d</u>ouble-<u>s</u>tranded DNA (dsDNA) templates (4). Precise genome edits are reported to be generated at low rates compared to standard indels and must be discriminated from a mixture of other outcomes, namely standard NHEJ-induced indels, <u>w</u>ild-<u>t</u>ype (WT) sequence and alleles with imprecise incorporations of template sequence. This low rate and diversity of outcomes presents challenges in 1) the analysis of injected F_0_ embryos to determine whether a particular gRNA/repair template combination shows promise and 2) the identification of germ-line transmitting F_0_ adults.

Here we have focused on precise-genome editing using ssODNs to create single amino-acid changes in target genes, which has been proposed to occur via <u>h</u>omology-<u>d</u>irected <u>r</u>epair (HDR) (2). Two attractive aspects of the ssODN approach are 1) the relatively short length (<150 nt) of the ssODNs, which can be commercially synthesized to eliminate in-house cloning or *in vitro* synthesis steps and 2) the correspondingly short genomic target that needs to be sequence-verified for potential divergence from the reference genome sequence. An arguable disadvantage of the ssODN approach is the lack of a fluorescent reporter gene, utilized in various dsDNA plasmid-based approaches, notably that by Hoshijima *et al.*, which allows for *in vivo* screening to detect germ-line transmission from F_0_ founders to F_1_ progeny (5). By contrast, screening when using the ssODN approach requires *ex-vivo* tissue samples being subjected to either direct sequencing or detection of gain or loss of a RE recognition site that was introduced by the repair template.

For standard CRISPR/Cas9 indel generation, we have learned through experience that selection of winning gRNA/Cas9 combinations is vastly aided by the ability to quickly estimate the frequency of targeted events in injected F_0_ embryos. Our method of choice for this has been the CRISPR <u>s</u>omatic <u>t</u>issue <u>a</u>ctivity <u>t</u>est (CRISPR-STAT), which measures the frequency of targeted indel formation in single embryos by combining fluorescent PCR and capillary electrophoresis (6). Here we describe our development of a CRISPR-STAT extension, tailored to estimate the frequency of genomic incorporation of RE recognition sites included in the ssODN template via synonymous substitutions, a method we have named TIARS (<u>t</u>est for <u>i</u>ncorporation of <u>a</u>dded <u>r</u>ecognition <u>s</u>ites).

The concept of TIARS emerged from our efforts to modify the *atp7a* gene in zebrafish. Loss of the human X-linked gene *ATP7A* causes Menkes disease, a disorder of copper transport that leads to death during the first few years of life if untreated (7,8). Consistent with copper’s essential role in pigment formation and angiogenesis, sequelae of Menkes disease include hypopigmentation and vascular abnormalities, and a current life-extending intervention combines early diagnosis with subcutaneous copper injections. The zebrafish Atp7a protein is 77% similar to the human ATP7A protein, including short stretches of near- or complete identity where cognate amino acids (AAs) are readily found. Although X-linked deficiencies cannot be modeled in zebrafish due to their lack of chromosomal sex determination, homozygosity of a null *atp7a* allele (named the *calamity* mutation, also called *atp7a^vu69^*) causes albinism in zebrafish larvae and death around 6 days post-fertilization (dpf) (9). In addition to Menkes disease arising from complete loss of *ATP7A*, a distinct human syndrome has been described for individuals carrying T994I or P1386S missense mutations in *ATP7A:* an isolated, adult-onset distal motor neuropathy (10–12). We initially developed TIARS to support our project to generate analogous missense alleles in zebrafish for future functional studies, *i.e.*, to quickly monitor our success at combining Cas9, gRNA and ssODNs to induce AA coding modifications in *atp7a* at the cognate zebrafish positions of T979I and P1387S.

Here we report 1) our use of TIARS to detect generation of both these alleles at the somatic level, 2) germ-line transmission of the *atp7a^T979I^* allele, which causes a pigmentation deficit in homozygous larvae and adults, 3) development of software to automate design of ssODN donor templates optimized for TIARS and 4) validation of this software by A) running it against a set of theoretical human targets and B) by high-throughput sequence analysis outcomes in the generation of a third zebrafish allele (the *ryr1b* gene).

## MATERIAL AND METHODS

### Zebrafish husbandry

EK strain wild-type (WT) zebrafish were used for most of this study, except for the high-throughput sequencing experiments, which used Tab5 strain WT zebrafish. Zebrafish were maintained in the aquatic animal facility under an Animal Use Protocol approved by the Institutional Animal Care and Use Committee (IACUC) of the *Eunice Kennedy Shriver* National Institute of Child and Human Development, MD, USA. All zebrafish embryos & larvae were maintained in egg water (0.006% sea salts and 0.1% methylene blue) in 10 cm Petri dishes at 28.5° C for a maximum of 7 days. Larvae selected for further growth were transferred to tanks and fed no later than 6 days post fertilization (dpf).

### Design and synthesis of sgRNAs

Protein sequence alignment identified T979 and P1387 of zebrafish Atp7a as cognate amino acids (AAs) of T994 and P1386S of human ATP7A. Zebrafish exon 15 for T979 and exon 22 for P1387 containing the AA codons to be modified from T and P to I and S, respectively, were analyzed on Ensembl. Candidate gRNAs were identified near the target mutation sites using the NHGRI-1 (ZebrafishGenomics) Track Data Hub on the UCSC genome browser (ZV9/danRer7 assembly) and compared to more recent assemblies of the zebrafish genome with BLAST to identify any off-target matches of concern. Gene specific Alt-R CRISPR-Cas9 crRNAs (20 mers) were purchased from IDT and co-annealed with IDT’s tracrRNA (57mer) at 95°C for 5 min and gradually cooled to room temperature to form a functional gRNA duplex, according to the manufacturer’s protocol (IDT, Skokie, Il).

### Single-stranded oligo mutation donor (ssODN)

Asymmetric ssODNs (107 nt) complementary to the gRNA nontarget strands per Richardson *et al*. were designed (13). They contain the desired non-synonymous codon mutation plus additional synonymous mutations to eliminate the PAM site and introduce two restriction enzyme (RE) sites. The two synonymous nt substitutions for creating new RE sites were found with the aid of *re site finder* (http://resitefinder.appspot.com/about) and where more than one option was available, the more frequent *Danio rerio* codon usage was favored according to the Kazusa Codon Usage Database (https://www.kazusa.or.jp/codon/). Putative alterations were also screened to ensure they do not affect predicted exonic splicing enhancers (ESEs) using the RESCUE-ESE Web Server http://hollywood.mit.edu/burgelab/rescue-ese/ and to ensure they do not create or eliminate any predicted microRNA targets using an additional web site (http://mamsap.it.deakin.edu.au/~amitkuma/mirna_targetsnew/sequence.html) that no longer appears to be supported at the time of this submission. PAGE-purified ultramer ssODNs were purchased from IDT. The ssODN sequence for *atp7a^T979I^* is indicated in Fig. 1, the ssODN sequence used for *atp7a*^P1387S^ was (CATTTTAGCAGGAGGGAGGACAGCACTACAGACACAGAAGATAGAGCCATAGCAGCAGAGCCCATCCAGCTCTGAAGGACCAACCCCACAGGCATAAACACAC) and the ssODN sequence for *ryr1b^I4936T^* was (TGTTTCAGTGTTATCTGTTCCATATGTATGTGGGTGTAAGGGCCGGAGGAGGAACTGGAGATGAGATTGAAGATCCAGCTGGAGA)

**Figure 1.**
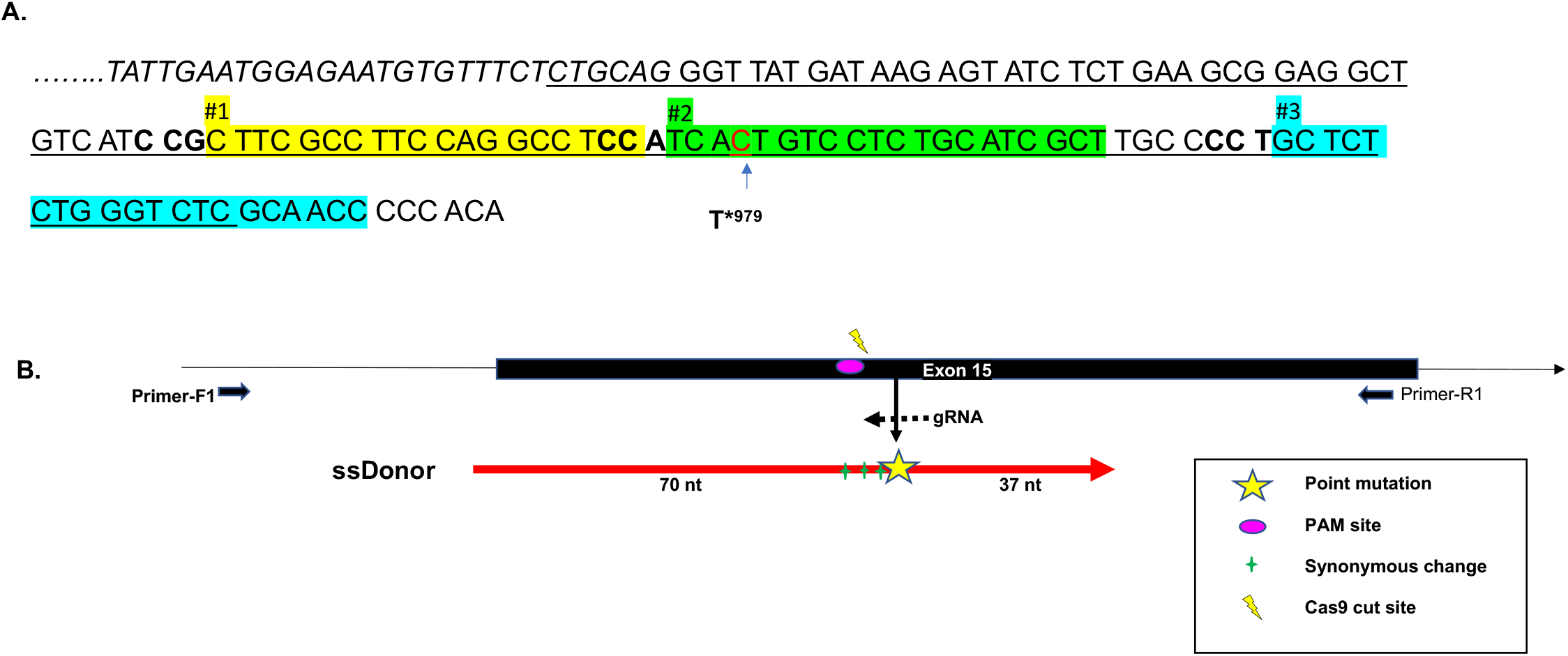
Knock-in strategy for generating the *atp7a^T979I^* allele. A. Genomic sequence of the zebrafish *atp7a* locus (exon 15) with the upstream intron (14–15) sequence italicized. Sequences corresponding to candidate gRNAs, each on the opposite strand, are highlighted (#1 in yellow, #2 in green and #3 in teal), and neighboring (opposite strand) PAM sequences are in bold. The #2 gRNA was selected for the editing project. The blue arrow indicates the T979 target codon and the full genomic sequence corresponding to the selected ssODN is underlined. B. Schematic representation of the knock-in and detection strategy. Atp7a exon 15, black box. The ssODN donor template (red) has the identical sequence as the targeted strand (*i.e.*, the reverse complement of the [sense] sequences shown in this figure) with the exception of four key changes (vertical arrows in detail), namely: one non-synonymous change (yellow star) and three synonymous changes (green stars) that together eliminate the PAM site (pink oval in schematic) and introduce ClaI and MluCI sites. Yellow lightning bolt on schematic indicates the double-strand cut site by Cas9 as directed by the gRNA (horizontal dashed arrow). Location of primers for detection (see Fig. 2) indicated as horizontal arrows. Note that primers flank all sequence corresponding to the single strand oligo donor, reducing the risk of amplifying the ssODN sequence from any other location or context.

### Microinjection

Cas9 protein (PNA Bio, Inc.) was mixed to a final concentration of 700 ng/ul with sgRNA (37.5 ng/ul) and ssODN (60 ng/ul), preincubated at 37°C for 5 min, centrifuged at 16,100 g 5 min to remove any debris and 1.4 nl was injected into the yolks of 1-4 cell embryos using the Pneumatic PicoPump system.

### Preparation of embryonic and adult fin lysates for genotyping

Euthanized 5 dpf embryos or fin biopsies from adult fish were treated with proteinase K (0.5 mg/ml) in 50 μl lysis buffer (20 Mm Tris-HCl (ph 8.9), 50 Mm KCL, 0.3% Tween-20 and 0.3% NP-40) at 55°C for 2 hr, 99°C for 5min and cooled to 4°C. Diluted lysate (1:10) was used as template for genotyping with primer sets flanking the sequence of the *atp7a* gene corresponding to the 107 nt of the ssODN.

### Primer design and fluorescent PCR assays

Two sets of primers [T979I editing-Forward: AGCAGTGAATGGGGAAGATG (Intron 14) and Reverse: TCCTCCCTTGATGAGGATTC (exon 15); and P1387S editing-Forward TGTTTGTCGTTTTAGTGAACC (intron 21) and reverse AAATAACCCAAATTATCATAACGGTCAC (intron 22)] were designed using Primer3Plus software to amplify 320 and 324 bp fragments, respectively, with the target AA roughly in the middle of the amplicon. M13 (TGTAAAACGACGGCCAGT) and PIGtail (GTGTCTT) adapters sequences were added to forward and reverse primers respectively(6). PCR reactions contained 1 μl of 1:10 diluted lysate and 5 μl premixed AmpliTaq-Gold (0.3U, Life Technologies) with 0.26 μM of target-specific- and an additional fluorescent forward primer: 6FAM-M13F (6FAM-TGTAAAACGACGGCCAGT)(6). In modified fluorescent PCR fragment analyses, the 6-FAM was directly linked to a forward primer with no M13 adapter sequence, and the 6FAM-M13F primer was not included. Amplification was performed on a MasterCycler, (Nexus gradient, Eppendorf) with initial denaturation at 95°C for 12 min, followed by 35 cycles of (95°C/30s, 57°C/30 s, 72°C/30s), and a final extension at 72°C for 10 min followed by a hold at 4°C.

### Screening for site-specific mutations

Restriction fragment length polymorphism (RFLP) analysis after the fluorescent PCR reaction was used to discriminate between wild type (WT) and point mutation knock-in alleles in F_0_ mosaic carriers and their F_1_ progeny. PCR products (5 or 10μl) were digested in the absence (control) or presence of designated REs in a total volume of 15 μ for 8h at 37°C. Samples (2μl) were then mixed with 10μl of Gene Scan 400HD ROX dye standard (1:50) in HiDi-formamide (Life Technologies) followed by denaturation at 95°C for 5 min and a hold at 4°C. RFLP fragments were separated in a genetic analyzer (ABI model 3130) using POP-7 polymer. Injection times were varied (23, 60 or 180s) according to experimental design. The default injection time was was 23 seconds. Data files (.fsa) were analyzed using GeneMapper or PeakScanner software (Thermofisher).

### Sequencing

For Sanger sequencing, PCR products were first treated with Exo-SapIT (Affymetrix) to eliminate unincorporated primers. Sequencing was outsourced (Macrogen USA) either (1) directly after PCR or (2) for early attempts at *atp7a*^T979I^ editing, after colony PCR using a nested reverse primer (ACTTGGTCAGGTTTCTATTTTG). Sequencing data was analyzed by Sequencher version 5.4.6. For high-throughput sequencing of *ryr1b* amplicons, injected 3 dpf embryos were lysed in groups of 10 and 1:10 dilutions were used as templates for PCR. Primers (F target-specific sequence: GCCTGACTGAGATCATGTCTTC; R target-specific sequence: ACCACACGGTAAAGTTCATACTCG) immediately flanked sequence corresponding to the ssODN, with no overlap, and included 5’ Illumina adapter sequences (F adapter: TCGTCGGCAGCGTCAGATGTGTATAAGAGACAG; R adapter GTCTCGTGGGCTCGGAGATGTGTATAAGAGACAG). PCR amplicons (198 nt in size) were cleaned using the Qiagen QIAquick PCR purification kit according to the manufacturer’s protocol and concentrations were assessed by Qubit and mixed at equimolar ratios with an amplicon distinct from *ryr1b* (unpublished data). Input concentration of mixed amplicons for second-round PCR-based library construction ranged from 3 – 10 ng/ μl. Each library was given a unique bar code (Nextera XT library prep kit), purified, pooled with other bar-coded libraries and subjected to 2 x 150 bp paired-end sequencing on an Illumina MiSeq system using the MiSeq Micro v2 Reagent Kit. Demultiplexing was performed by bcl2fastq v2.20.

### Computer assisted ssODN template design

TIARS <u>Designer, described below, is an open-source Python package available at</u> https://github.com/NICHD-BSPC/tiars-designer.

## RESULTS

### Generation of *atp7a^T979I^* and *atp7a*^P1387S^ variants

We sought to create zebrafish models for human distal motor neuropathy caused by human mutations cognate to hypothetical zebrafish mutations atp7a^T979I^ and *atp7a^P1387S^*. Reports on ssODN-based CRISPR/Cas9 precise editing in zebrafish and other systems indicate that two key criteria driving efficiency are 1) minimizing the distance between the cut site (situated between position −3 and −4 5’ to the PAM NGG) and the targeted edit, and 2) maximizing the cutting efficiency of the gRNA (3,14). For the *atp7a^T979I^* edit (a C>T change altering the codon threonine/ACT to isoleucine/ATT; Fig. 1B), we considered the three closest PAM-adjacent targets to the *atp7a^T979I^* edit site, each on the antisense strand and numbered according to their 5’ to 3’ locations: GGAGGCCTGGAAGGCGAAG (#1, yellow), AGCGATGCAGAGGACAGTGA (#2, green) GGTTGCGAGACCCAGAGAGC (#3, teal). The ranking of predicted distances between the edit site and cut sites for these three candidates is: 0 <u>n</u>ucleo<u>t</u>ides (nt; #2) < 20 nt (#1) < 25 nt (#3). To determine cutting efficiencies of the gRNAs corresponding to these three targets, we used CRISPR-STAT as follows. gRNAs corresponding to *atp7a^T979I^* targets 1-3 were individually co-injected into zebrafish embryos with recombinant Cas9 and individual embryos were harvested for DNA several days later. The targeted locus was PCR amplified with a primer mix that included a fluorescent label, and amplified fragments were subjected to capillary electrophoresis to reveal their migration and fluorescent intensity of each labeled PCR fragment. Cutting efficiency was calculated as the sum of non-WT peak intensities divided by the sum of all peak intensities (data not shown). Cutting scores of the three targets ranked as follows: #1 (97%) > #3 (30%) > #2 (7%) (Table S1). Target #2 had the best proximity but lowest cutting score. We considered proximity to be the more important characteristic in this case because gRNAs with very high cutting scores can cause extensive somatic cell mutation. In essential genes such as *atp7a*, such widespread mutations are incompatible with survival and establishment of germ-line transmitting founders. Indeed, the gRNA against target #1 fully recapitulated the lethal *calamity* null *atp7a* phenotype, including death by 6 dpf (data not shown), and so we were regardless forced to consider scenarios of lower cutting levels.

The single strand donor template (ssODN) for precise editing was designed following the rules suggested by Richardson *et al.*, namely with sequence complementary to the non-target strand of gRNA#2 and with homology arms of unequal length flanking the desired modification with a bias to the 5’ side (13). Thus, we selected (5’) 70- and (3’) 37- nt homology arms (Fig. 1A underlined & 1B). In addition to including the targeted non-synonymous C>T change, we made three synonymous changes. First, following published recommendations aimed at reducing the chance of re-targeting and re-cutting of successful edits, we eliminated the PAM site for target #2 with a C>G change (3). Second, we introduced two additional synonymous C>A changes to create novel <u>RE</u> sites for ClaI and MluCI, in order to facilitate downstream F_0_ and F_1_ screening. The C>T missense mutation underlying the T979I change was included in the MluCI RE site.

For *atp7a^P1387S^* variant generation we used analogous reasoning and strategies, selecting a gRNA candidate with a 24% cutting score (Fig. S1) and cutting within 5 nt of the targeted edit (Fig. S1). In addition to the central non-synonymous CC>AG changes (CCC/proline to AGC/serine), we introduced two additional synonymous changes (T>C and G>T) creating novel RE sites for HpyAV and Hpy188I. Analogous to MluCI for *atp7a^T979I^*, Hpy1881 recognizes the C>A change that is part of the P1387S change. As a further convenience, there is a nearby endogenous Hpy1881 that could serve as an internal control for Hpy1881 activity. Because of the PAM site’s registration within the coding sequence, there was no way to disrupt it with synonymous changes. Thus, we were able to find potential gRNAs and RE-site generation strategies for each of the alleles we wished to create.

### Screening for targeted *atp7a* edits and development of TIARS

Because of the multiplicity of mutagenic outcomes mosaically represented among somatic cells of injected F_0_ embryos, direct Sanger sequencing is not a viable strategy for assessing precise editing rates. Our initial strategy for assessing the rate of *atp7a^T979I^* edits was instead to lyse pools of injected F_0_ embryos (n=10, 5 dpf), PCR amplify the targeted region, and then subclone and sequence colonies. Out of 96 colonies analyzed, three had precise edits, 44 had unaltered WT sequence and 49 had various indels. We suspect, however, that these three precise edits represented false-positive artefacts. This is based on the following observations. First, further screening of 288 individual F_1_ progeny from 16 F_0_ founders revealed template insertions only. Second, subcloning from one of these insert-carrying F_1_ progeny revealed that the subcloning process itself can convert tandem repeats into precise edits, presumably due to bacterial recombination, as has been described (Fig. S2) (15).

As an alternative to subcloning, we developed TIARS by modifying CRISPR-STAT as follows. After the fluorescent PCR step, but prior to capillary electrophoresis, a portion of the PCR product is RE digested to visualize the <u>r</u>estriction <u>f</u>ragment <u>l</u>ength <u>p</u>olymorphisms (RFLP) associated with precise edits that might be mosaically present in individual F_0_ embryos. Our initial TIARS protocol used the cost-saving three-primer strategy of CRISPR-STAT, comprised of a “universal” 5’ FAM-labeled M13 primer, an M13-tailed forward primer and a reverse primer (Fig. 2) (6). Two of 44 F_0_ embryos 5 dpf after microinjecting gRNA/ssODN/Cas9 protein showed a low fluorescent RFLP signal (Fig. 2A) with the correct size (+/− 1 nt) of the predicted ClaI and MluCI fragments. However, because the intensity of these signals was not far above background noise, we sought to improve the TIARS signal intensity by replacing the three-primer strategy with a two-primer strategy in which the forward primer itself carries the 5’ FAM fluorophore, ensuring quantitative labeling of the amplicon. Comparing the two approaches, the fluorescent RFLP signals increased from fewer than 40 fluorescent units (FLU; Fig. 2A) to over 400 FLU (Fig. 2B). Under this modification of TIARS, we found 2 of 31 F_0_ adults carrying the correct integration in fin biopsies. All told, we observed a somatic editing rate of 4 out of 75 (5.3%) F_0_ zebrafish for the *atp7a^T979I^* edit. Beyond direct labeling of the forward primer, we further augmented sensitivity by increasing the quantity of PCR product loaded onto the ABI machine and the programmed injection time, the latter of which determines the fraction of sample actually loaded onto the electrophoretic capillary (Fig. S3).

**Figure 2.**
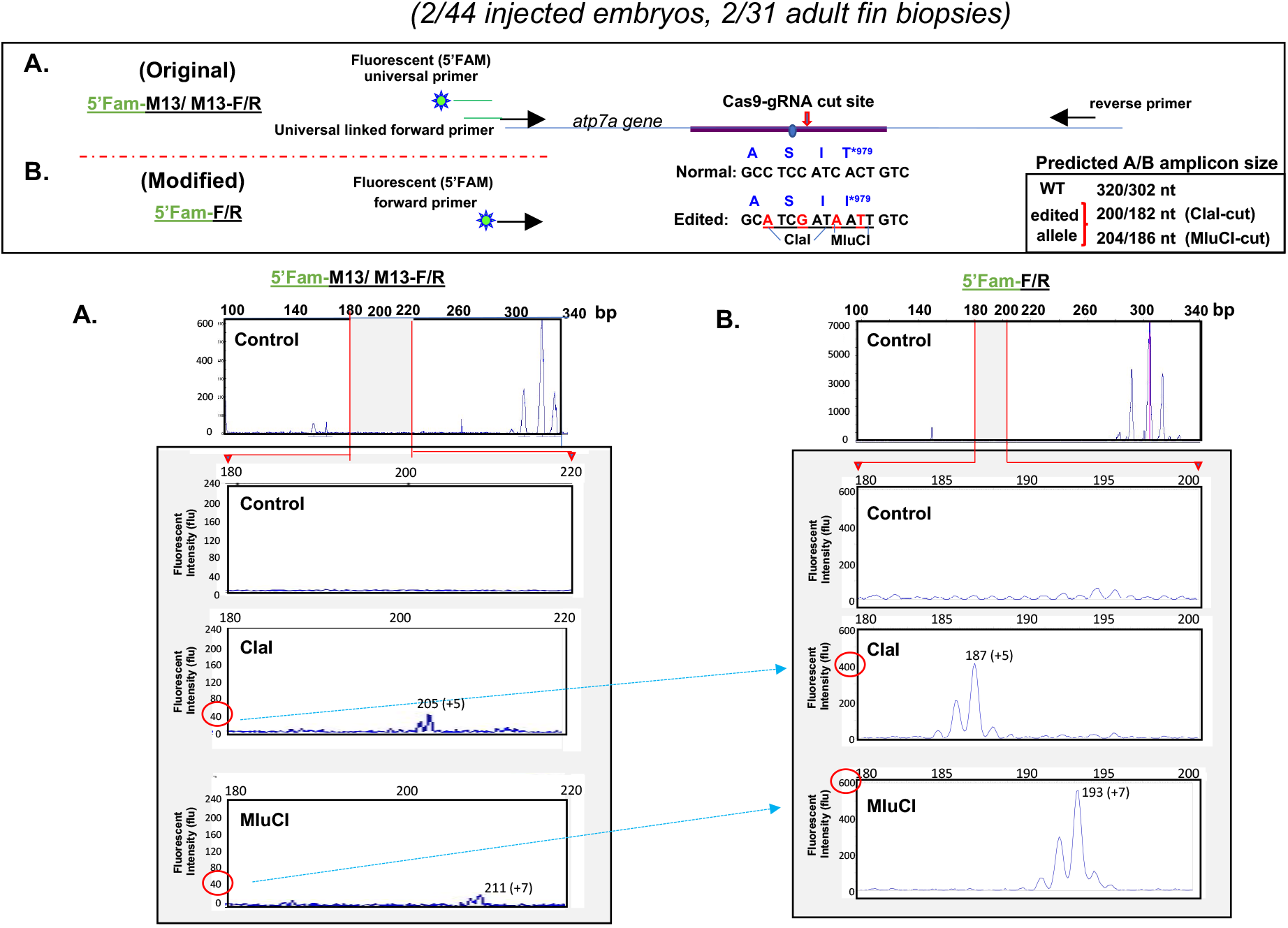
F_0_ screening strategy for ssODN-directed changes by RE digestion of fluorescent PCR prior to fragment analysis. Top panel: Schematic representation of original (A) versus modified (B) fluorescent PCR fragment analysis. Original approach of PCR reaction with three primers, 5’FAM-M13 and M13 tailed Forward and Reverse primers. Modified PCR reaction with two primers, 5’FAM directly linked to the forward primer and reverse primer. Predicted amplicon sizes using 3-primer/or 2-primer strategies (18 nt difference due to presence/absence of M13 sequence on 6FAM-labeled strand) are indicated in the corner box. Lower panel: Fluorescent PCR was performed using the original method (A) and modified approaches (B). Results are shown are for an adult fin biopsy. The PCR product was digested in the absence (Control) or presence of REs ClaI or MluCI prior to analysis. Migration distance and flluorescent intensity of the 5’ labeled strand spanning from the 6-FAM label to the other end of the amplicon or the 5’ RE cut site was measured on an ABI 3130 Capillary Electrophoresis unit. The uncut amplicon size was either A=320 nt or B=302 nt depending on the presence or absence of the 18 nt M13 sequence. A) Pre-digestion with ClaI or MluCI led to the appearance of novel low-intensity peaks below 40 fluorescent units (flu) close to the predicted sizes for a precisely edited allele. We attribute the discrepancies of +5 nt for the ClaI fragment (205 nt observed vs. 200 nt predicted) and +7 nt for the MluCI fragment (211 observed vs. 204 nt predicted) to sequence-dependent variance of fragments from molecular standards, as frequently observed on the ABI 3130, with relatively larger effects on relatively smaller fragments. B) Direct labeling of the 5’ primer increased sensitivity more than 10-fold, with the novel peaks now showing intensities >400 flu (compare A and B). No other REdependent peaks of this magnitude were observed. As with the low-intensity peaks in A), there was a variance of +5 nt (187 nt observed vs. 182 predicted) and +7 nt (193 nt observed vs. 186 nt predicted), all observations and predictions being reduced by 18 nt, in accordance with the elimination of the 18 nt M13 sequence from the modified 6FAM-labeled 5’ primer.

TIARS on embryos injected to create *atp7a^P1387S^* revealed 4 out of 41 F_0s_ (9.8%) with the predicted diagnostic pattern for Hpy1881 and HpyAV (Fig. S4). We used this same scenario to address whether phosphorothioate (PS) end-protection of the ssODN repair template might increase the rate of precise editing, as has been reported (16). However, in our hands, co-injecting a PS-protected version of the repair template yielded a lower incidence of F_0s_ with the predicted diagnostic pattern: 3 out of 81 (3.8%; Fig. S4). Unfortunately, no F_0_ embryos survived from these standard ssODN and PS-ssODN experiments, likely because the gRNA was too effective and replicated the lethal phenotype of *atp7a/calamity* loss of function. In summary, we successfully established TIARS as a screening tool and identified two potential F_0_ carriers of the *atp7a^T979I^* mutation.

### Germ-line transmission of the *atp7a^T979I^* allele

Our screen for germ line transmission was straightforward, as we successfully crossed a WT fish with one of the two F_0_ animals that had had a diagnostic TIARS signal from its fin biopsy, and one out of the 32 F_1_ progeny obtained was found to be heterozygous for the modified allele according to TIARS (Fig. 3A and B), the integrity of which was then confirmed by sequencing (Fig. 3C). Per zebrafish research community convention, we gave this novel *atp7a^T979I^* allele the alternate name of *atp7a^y652^*, the “y” indicating its institutional origin (NIH/NICHD). Applying Human Genome Variation Society rules to zebrafish, with the assistance of the Mutalyzer website (https://mutalyzer.nl), we also determined the following non-ambiguous full name for this allele: NM_001042720.1(atp7a):c.2928_2936delCTCCATCACinsATCGATAAT. For TIARS on F_1s_, we returned to our initial 3-primer version of the protocol. This was because the sensitivity required for detecting rare cells in F_0s_ is in fact too high when genotyping embryos that carry the same genotype in every cell. We crossed this heterozygous *atp7a^y652/+^* animal with a heterozygous carrier of the *calamity* allele, *atp7a^vu69/+^*, and scored and genotyped progeny, revealing a subtle hypopigmention phenotype of *atp7a^y652/vu69^* trans-heterozygous embryos (Fig. 3D-E), a phenotype that persists into adulthood (Fig. 3F-G), confirmed both by Mendelian ratios and by genotyping. Thus, with the help of TIARS, we have established a novel model for zebrafish *atp7a* dysfunction. Research on the general impacts of the *atp7a^y652^* allele on zebrafish health and physiology are ongoing and will be reported elsewhere.

**Figure 3.**
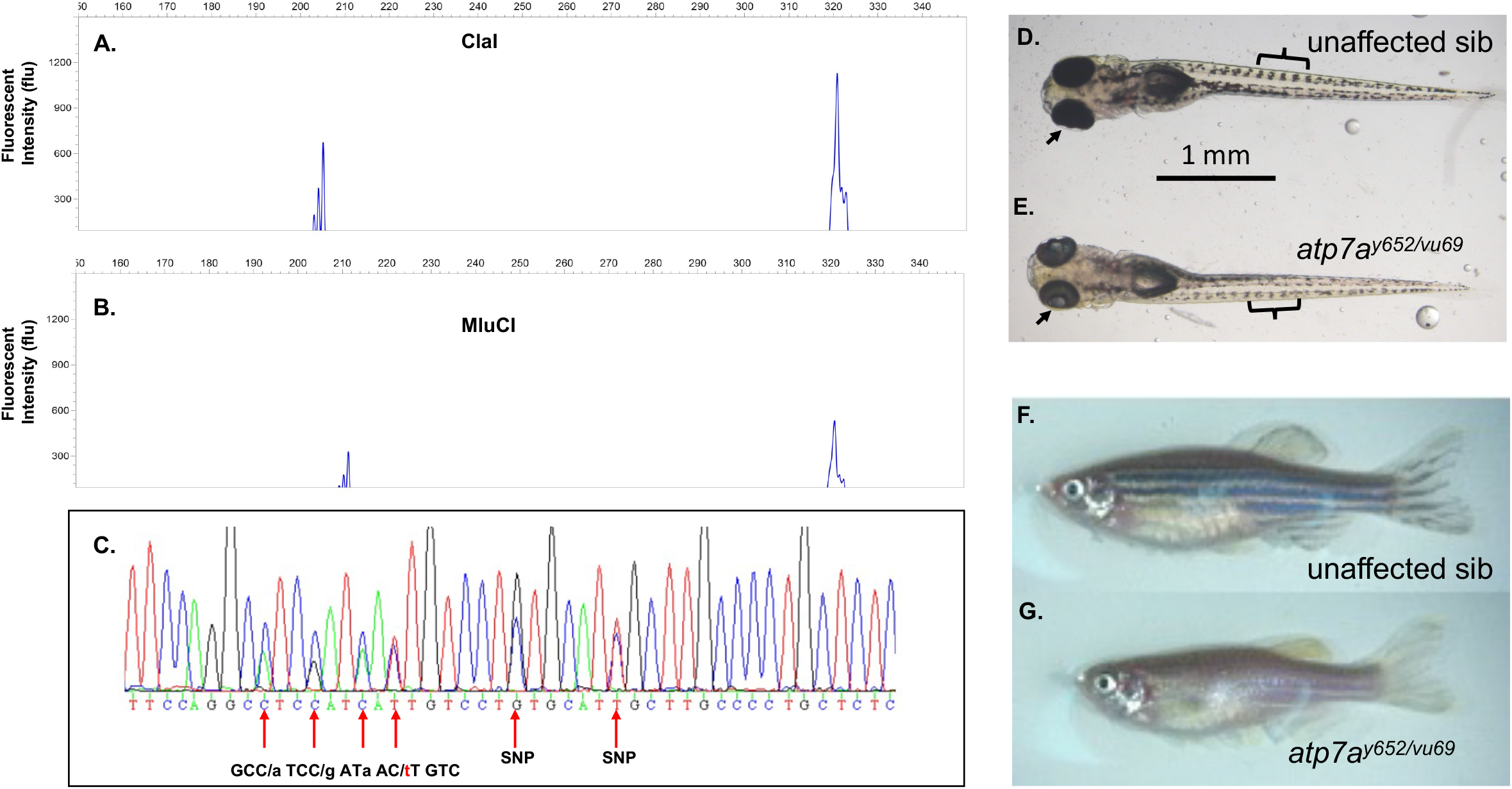
Germ-line transmission and sequence verification of the *atp7a*^T979I^ edit. **A-B**. Germline transmission was demonstrated by heterozygosity of the allele of interest among F_1_ embryo progeny from an F_0_ female candidate carrier (based on previous fin biopsy results as per Fig. 2) crossed with a WT male. ABI electrophoretic traces showing wild-type alleles at 320 nt and precise-edited allele after RE digestion with ClaI at 205 nt (200 predicted) and MluCI at 211 nt (204 nt predicted). **C**. Sequence verification. Arrows indicate mixed WT/edited-allele peaks at each of the four targeted positions, as well as mixed peaks at two single-nucleotide polymorphisms (SNPs) known to be present in the EK zebrafish strain. **D-G**. Hypopigmented phenotype of *atp7a^y652/vu69^* larval and adult fish. **D-E**. Representatives of the two phenotypes seen among 5 dpf progeny of a cross between an *atp7a^y652/+^* female and an *atp7a^vu69/+^* male. 75.7% (143 of 189) had the standard WT phenotype (D), characterized by very dark eyes (arrow) and melanocytes (bracket). 24.3% had a subtle but discernibly hypopigmented phenotype (E) with lighter eyes (arrow) and melanocytes (bracket). Subsets of these dark and light fish were raised in separate tanks and the hypopigmented phenotype persisted (compare G with F) at 3 months postfertilization. Genotyping of fin biopsies from the hypopigmented cohort revealed that 100% (16/16: χ^2^ P = 1.2 x 10^−5^) had the *atp7a^y652/vu69^* genotype, whereas fin biopsies from the dark cohort yielded representatives of the other three genotypes, namely *atp7a^vu69/+^* (5/22)*, atp7a^y652/+^* (8/22) and *atp7a^+/+^* (9/22), but none of the *atp7a^y652/vu69^* genotype (0/22: χ^2^ P = 1.2 x 10^−5^ = 0.012).

### TIARS Designer: a computer-assisted repair template design tool

Manual identification of suitable synonymous sequence alterations to generate novel RE sites is a time-consuming and error-prone process. To assist with future precise editing projects, we developed TIARS Designer, which exhaustively searches all possible RE site combinations, filters them to retain those that meet certain criteria, and ranks the remaining options for final manual selection. An overview of the computational steps is provided in Fig. 4. The inputs to the program are 1) approximately 200 nts centered on the PAM-adjacent target of interest, 2) the PAM-adjacent target sequence, and 3) the desired non-synonymous change.

**Figure 4.**
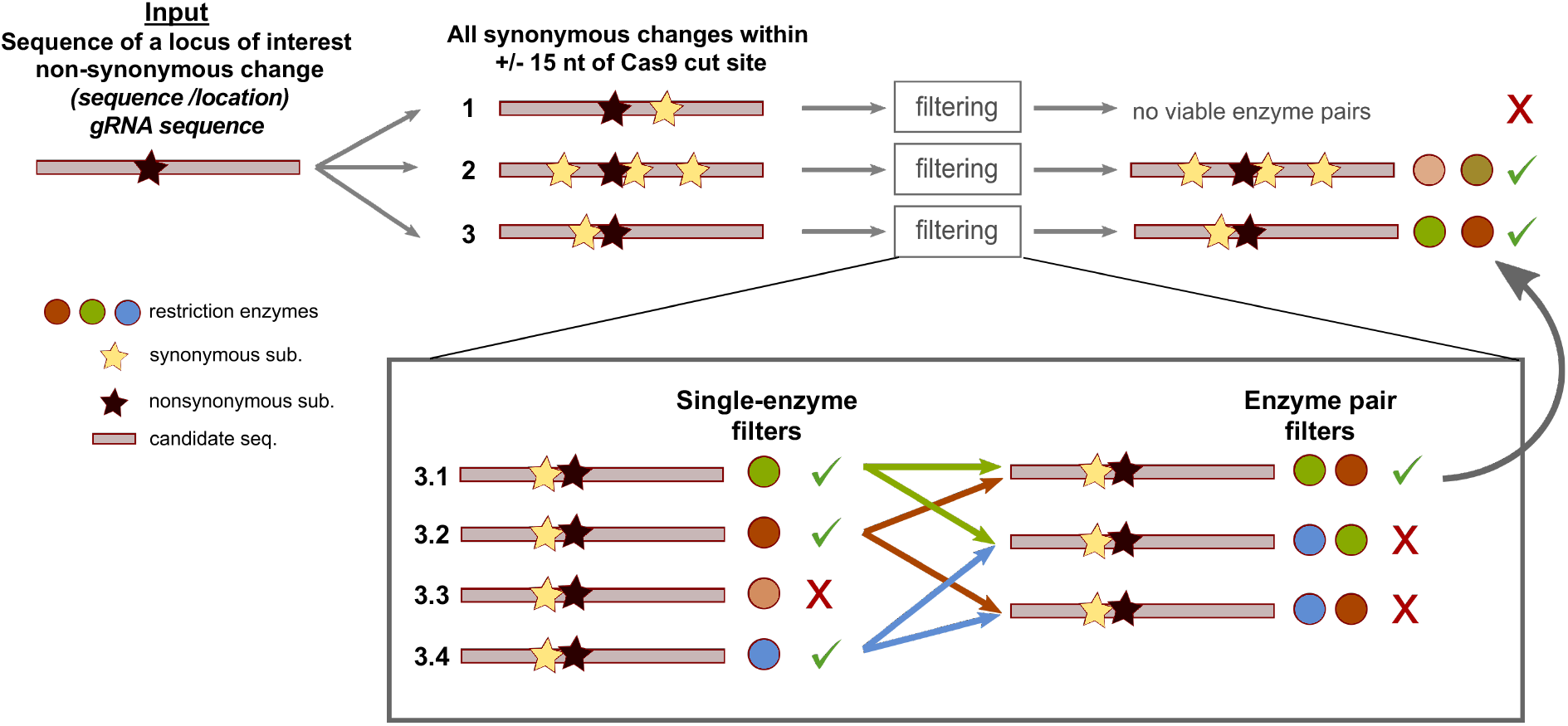
TIARS Designer flow chart. Repair template candidates are generated as follows. The data entered is the locus sequence with the desired non-synonymous change and the location of the predicted cut site (between position −3 and −4 of the NGG PAM for Cas9). The first computational step generates the set of sequences carrying all possible combinations of synonymous changes within 15 nt of the predicted cut site. Each of these sequences is then subjected to two sequential sets of pass-fail queries: the Single Enzyme filter and the Enzyme Pair filter, as detailed in Fig. 5.

In the first step, sequences spanning from 15 nt 5’ to 15 nt 3’ of the Cas9 cut site are mutated *in silico* to generate the set of all possible cut site-proximal synonymous substitutions (Fig. 4). We limited the *in silico* synonymous substitutions to +/-15 nt flanking the cut site for two reasons. First, we assume that users will in general only take on such a project if they have a PAM-adjacent sequence for which the cut-site is within 15 nt of the targeted change, as is generally recommended (3). Second, our ssODNs are typically in the 90-110 nt range, and we wished to conserve 100% homology in the more distal portions. Depending on the identity of the amino acids, the possible number of candidate sequences containing synonymous substitutions within 15 nt of the cut site could range from zero (in the unlikely case where all ten amino acids are methionine or tryptophan which only have a single codon) to 6^10^ (in the case where all ten amino acids are arginine with 6 possible codons).

The entire set of candidate modifications is then screened against a database of commercially available REs. Each combination of RE and candidate sequence must pass a set of filters (Fig. 5) to move on to the next stage. Filter I.1 ensures each novel RE site is created by at least two synonymous substitutions. This is to limit the noise of frequent NHEJ-induced single nucleotide changes creating a RE site, a phenomenon we observed in our first attempts at TIARS with different REs (data not shown). RE sites for which nearby endogenous sites exist are attractive because the endogenous sites can serve as positive controls for RE cutting and help avoid false-negative conclusions, but the orientation and spacing of these endogenous sites needs to be carefully considered. Filter I.2 requires that no endogenous cut site occur within five nucleotides of a novel cut site so that the two fragment sizes can be reliably discerned by capillary electrophoresis. Filter I.3 ensures no endogenous RE sites flank a novel RE site on both sides. Otherwise, when PCR amplicons are digested, the labeled fluorescent primer will be separated from the diagnostic cut, regardless of which side of the amplicon it is on. Finally, filter (I.4) ensures the number of RE recognition sites are equal to the number of RE cut sites for a given RE. This is to avoid scenarios where the RE recognition site is present but not the RE cut site, which can occur with Type II REs whose cut sites are outside of the recognition site. As an additional strategy against false positive hits, we built our approach around creating two unique RE sites. Therefore the next set of filters considers pairs of REs for each candidate sequence. To be included in a RE pair for a candidate sequence, both REs must have passed the above set of filters for the sequence.

**Figure 5.**
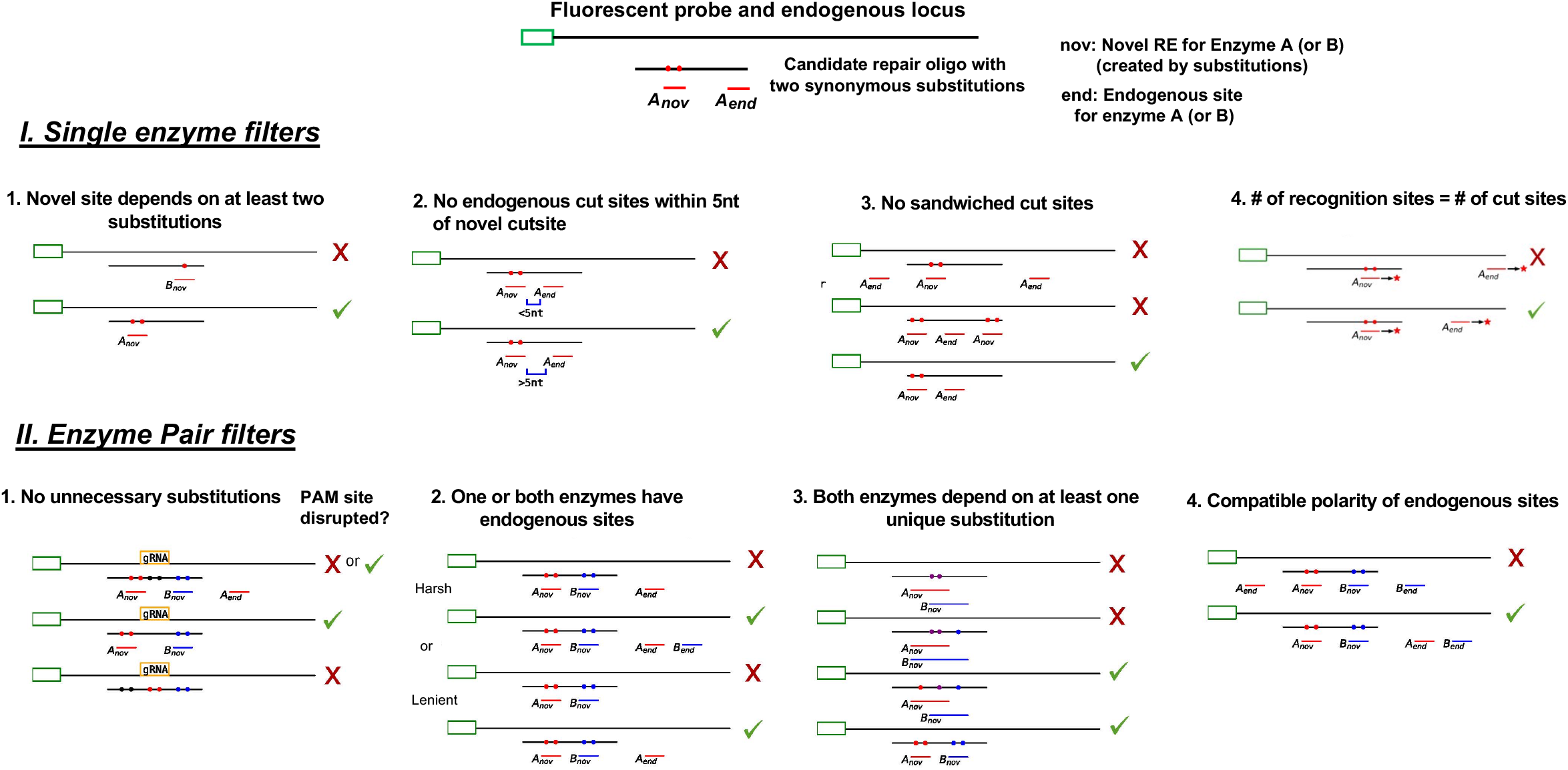
TIARS Designer filter criteria. Repair template candidates have to pass four tests each for the Single Enzyme filter (I) and the Enzyme Pair filter (II).

Filter II.1 ensures there are no unneeded substitutions. This avoids any superfluous nt substitutions, which in theory could diminish the rate of HDR. But an exception is made under Filter II.1 if the registration of the PAM site relative to the coding sequence is such that it cannot be disrupted by synonymous substitutions, leaving a risk of gRNA/Cas9 re-cutting of the altered allele. In these instances, a maximum of synonymous changes neutral to the novel RE design are introduced in the 10 nt 5’ to the PAM site, the region of the PAM-adjacent target that is thought to be most important for Cas9 targeting specificity, in the hopes of inoculating the altered PAM-adjacent target against gRNA/Cas9 re-cutting (17). Filter II.2 considers endogenous sites. It has two settings: the first (harsh) looks exclusively for pairs in which each novel RE also has an endogenous site. The scenario of each RE having an endogenous site is rarer, however, and so a second Filter II.2 setting (lenient) accepts pairs where only one novel RE has an endogenous site. Filter II.3 requires that novel RE recognition sites be formed from at least one unique substitution upon which the other novel site does not also depend. This ensures that the two REs act as proxies for distinct nt substitutions. Filter II.4 demands that endogenous sites of both novel REs be on the same side of the amplicon, a requirement that allows for the convenience and cost savings of having the fluorophore on the same side of the amplicon for TIARS of either enzyme. Any remaining pairs of REs and candidate sequence that pass these filters are then considered the final candidates. The output consists of Excel spreadsheets containing the sequence, the restriction enzymes, and various statistics recorded from the filters, such as distance from RE to the Cas9 cut site, distance from RE site to the amino acid change, and other information that could be useful in ranking the hits, like codon frequency metrics. In addition to this spreadsheet, the tool also outputs a Word document that uses typography to encode the various features in each candidate to help with manual inspection and making the final choice for a locus.

### TIARS Designer Examples and Validation

As a demonstration of TIARS Designer, Table 1 presents the numbers of candidates and solutions found for *atp7a^T979I^* and *atp7a^P1387S^* using the same PAM-adjacent targets that we selected and also for a new example: *ryr1b^I4936T^*, a zebrafish mutation that would be cognate to the human allele *RYR1^I4898T^*, which causes congenital myopathy (18). The enzyme pairs we had selected for *atp7a^T979I^* and *atp7a^P1387S^* were each one of 45 and 188 possible solutions, respectively, per TIARS Designer (Table 1). All solutions are presented as summary excel files and individual MS word documents, as shown for one of the *ryr1b^I4936T^* solutions (Fig. 6). The excel file includes the distance of the RE cut site to the targeted amino acid change and to the Cas9 cut site and the frequency of suggested codons in zebrafish. This information can be used to rank candidate solutions according to user concerns and preferences when multiple solutions are available.

**Figure 6.**
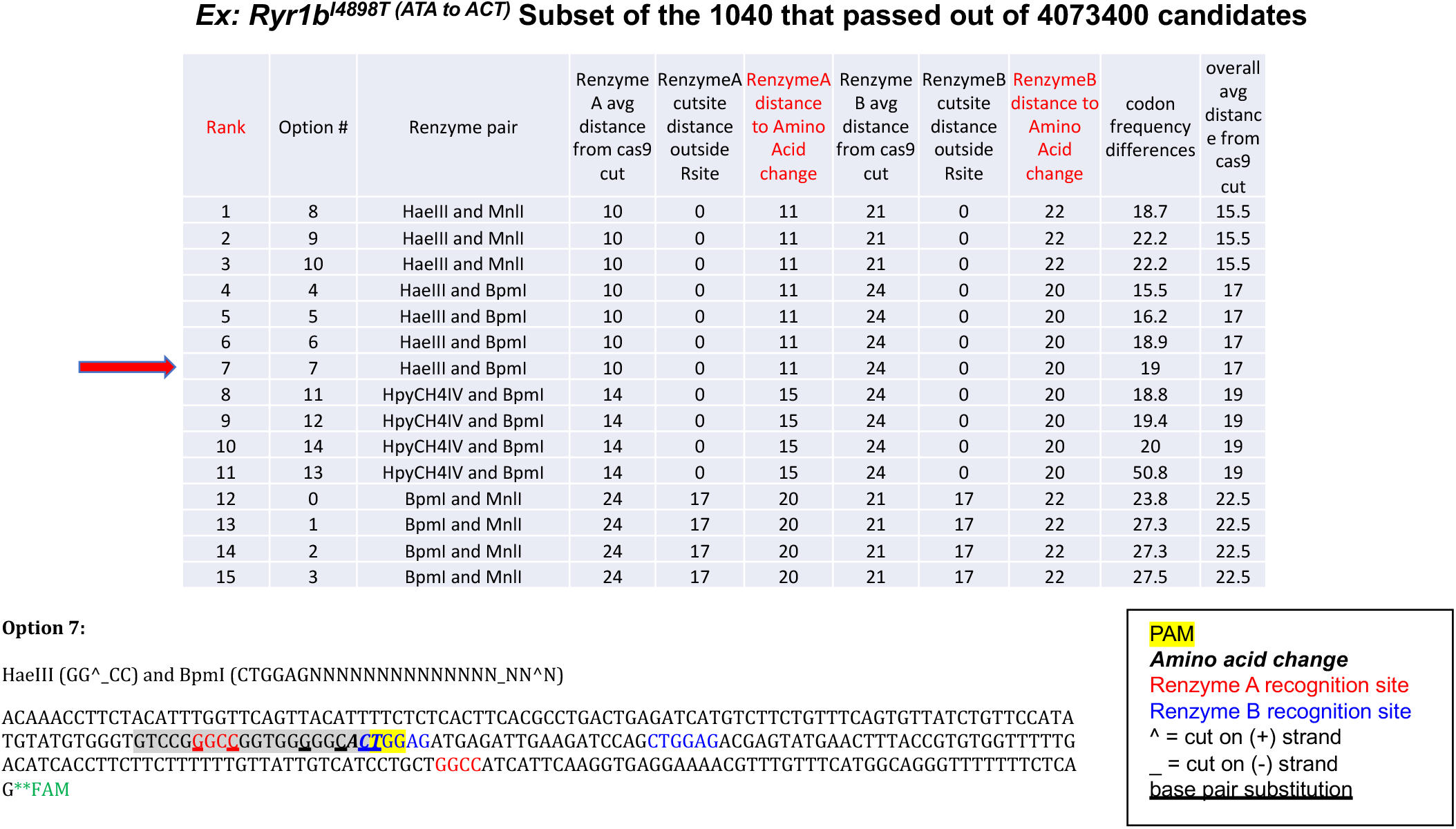
Sample outputs of TIARS Designer (example: *ryr1b^I4936T^*) Top: Summary list (Microsoft Excel) enabling a ranking of repair template candidates according to criteria provided in the column headers. Bottom: Individual pages (Microsoft Word) for each candidate with detailed annotations.

**Table 1.** TIARS Designer pass/fail statistics for the complete set of synonymous substitution candidates (within a 30 nt window centered on the cut site) for *atp7a^T979I^*, *atp7a*^P1387S^ and *ryr1b^I4936T^*.

| Sample | no sandwiched cutsites | no endogenous cutsites within Nnt of novel cutsite | novel site depends on at least two substitutions | num recognition sites equals num cutsites | no unneeded substitutions | Total Candidates before SE filtering | Total Candidates after SE filtering |
| --- | --- | --- | --- | --- | --- | --- | --- |
| **Atp7a P1368S** | 62.5 | 70.32 | 27.49 | 81.3 | 8.75 | 526344 | 10136 |
| **Atp1a T979I** | 60.2 | 66.27 | 25.12 | 70 | 0.06 | 498042 | 48 |
| **Ryr1b I4936T** | 71.08 | 76.43 | 37.47 | 84.86 | 14.1 | 5133504 | 212304 |

**Enzyme Pair Filters**
| Samples | at least one has endogenous site | both have endogenous site | compatible polarity | both Renzymes depend on at least one unique substitution | no unneeded substitutions | Total Candidates before EP filtering | Total Candidates after EP filtering |
| --- | --- | --- | --- | --- | --- | --- | --- |
| **Atp7a P1368S** | 37.68 | 2.36 | 98.06 | 47.94 | 19.54 | 158176 | 188 |
| **Atp7a T979I** | NA | NA | 100 | 45.8 | 0.16 | 38256 | 45 |
| **Ryr1b I4936T** | 13.76 | 1.21 | 99.54 | 58.4 | 18.43 | 4073400 | 1040 |

Continuing with this *ryr1b^I4936T^* example, we injected zebrafish embryos with an oligo containing these changes, pooled and lysed samples (n=10, 3 dpf), PCR amplified the targeted region and performed high-throughput sequencing as part of a larger study (manuscript under preparation). Analysis of the combined total reads from two such samples, as well as combined total reads from three negative controls in which Cas9 and the gRNA but no oligo were injected, indicates the following. First, that *ryr1b^I4936T^* has somatic rates of perfect edits representing 3.2% out the total reads and 16.6% out of those WT and perfectly edited reads that were not otherwise altered (6020 MUT reads out of 186870 Total Overall reads and 6020 MUT reads out of 39,262 MUT + WT reads, respectively; Fig. 7A “+” and Table 2.). The overall high rates of “OTHER” reads (Fig. 7A and Table 2), meaning reads that were altered in length or had non-synonymous changes beyond the targeted AA change, likely reflects a combination of mutagenesis due to erroneous NHEJ events, non-homologous inclusion of ssODN sequence and PCR/High-throughput sequencing-introduced errors, as has been observed in other high-throughput sequencing analyses of zebrafish HDR experiments (19). Second, our analysis indicates a 97.3% coincidence of reads (5850 of 6020 MUT reads) bearing the desired edit and the combination of novel REs and no other confounding changes to RE sites. This indicates that TIARS would theoretically have served as an excellent proxy for the desired edit (Fig. 7B and Table 2). Stated otherwise, there were only 2.7% candidate false-negative reads (170 of 6020 MUT reads) where TIARS would theoretically have failed due to a lack of diagnostic RE site creation or other changes to the diagnostic pattern. Also rare (0.06%; 19 out of 33,242 total WT reads) were falsepositive scenarios in which one or the other of the diagnostic novel RE sites was present but not the desired edit (Fig. 7C and Table 2). This low rate of false-positive outcomes indicates the requirement for hypothetical RE sites to be created from a minimum of two synonymous mutations (Fig. 5, filter 1.i) is sufficiently stringent.

**Fig. 7.**
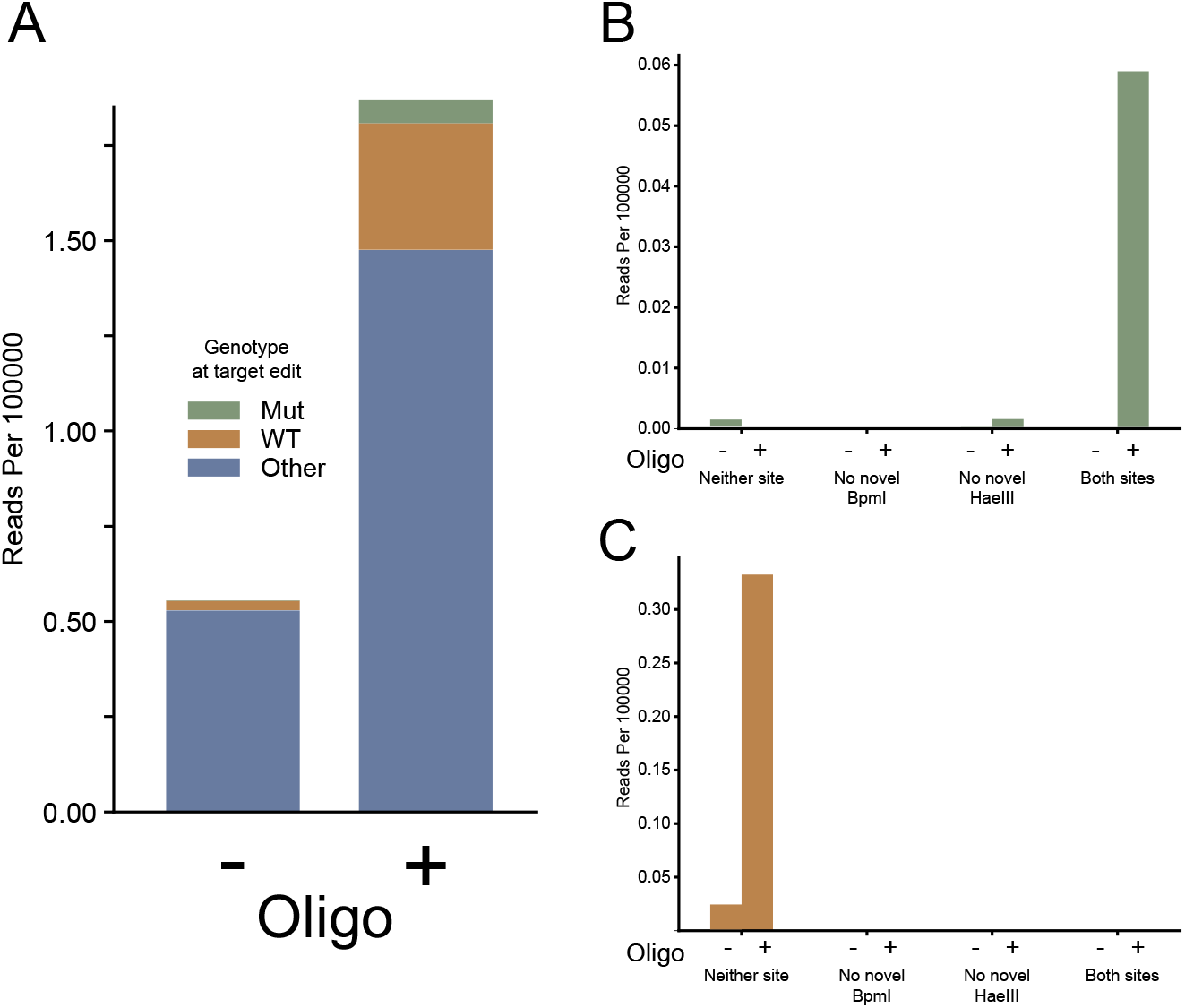
Sequence inventory of *ryr1b^I4936T^* edits and other outcomes among F_0_ (injected) embryos. Total reads from combined independent pools of ten 3 dpf Tab5 embryos each that had been injected at the 0-4 cell stage (0 dpf) with a combination of Cas9, the selected *ryr1b^I4936T^*-targeting gRNA (grey highlight, Fig. 6) and the ssODN (“+”, two independent pools) or without the ssODN (“-”, four independent pools). See Table 2 for further details.

**Table 2.** Inventory of high-throughput reads among F_0_ embryos targeted for creation of an *ryr1b^I4936T^* allele. Reads were initially classified as either encoding the unaltered WT AA sequence (WT), encoding the desired AA substitution only (MUT) or encoding additional AA changes (OTHER). Further classifications considered various combinations of the following features: presence (CC) vs. absence (no CC) of the targeted codon change; presence of both the novel BpmI and HaeIII RE sites (both Rsites)(All desired changes); absence of the endogenous BpmI site (no endog Rsite); presence of only one or the other of the novel BpmI or HaeIII sites (no novel Rsite)(only novel Rsite)(off target Rsite)(apparent novel Rsite); and absence of both novel BmpI or HaeIII sites (neither Rsite). The same categories were also applied and inventoried for the HpaII RE and an imaginary New Enzyme with RE sites that can be created by single nt substitutions, as opposed to the BpmI and HaeIII RE sites that require two nt substitutions in the context of *ryr1b^I4936T^*.

|  | **BpmI** | **HaeIII** | **HpaII** | **New Enzyme** |
| --- | --- | --- | --- | --- |
| **All desired changes (MUT)** | 5991 | 5879 | 5877 | 5862 |
| **CC\| no novel Rsite (MUT)** | 18 | 141 | 135 | 135 |
| **CC\| no endog Rsite (MUT)** | 5 | NA | NA | NA |
| **CC\| neither Rsite (MUT)** | 0 | NA | NA | NA |
| **no CC\| both Rsites (WT)** | 0 | NA | NA | NA |
| **no CC\| only novel Rsite (WT)** | 0 | 11 | 11 | 11 |
| **no CC\| neither Rsite (WT)** | 40 | NA | NA | NA |
| **CC\| off target Rsite (MUT)** | 6 | 0 | 8 | 23 |
| **no CC\| off target Rsite (WT)** | 8 | 0 | 64 | 58 |
| **off target CC\| apparent novel Rsite (OTHER)** | 0 | 0 | 0 | 0 |
| **category sum** | 6068 | 6031 | 6095 | 6089 |
| **total WT** | 33242 | 33242 | 33242 | 33242 |
| **total MUT** | 6020 | 6020 | 6020 | 6020 |
| **total OTHER** | 147608 | 147608 | 147608 | 147608 |
| **total overall** | 186870 | 186870 | 186870 | 186870 |

To dig in further, we next asked whether this “two or greater” synonymous mutations per novel RE site rule (Fig. 5, filter 1.i) is not only sufficient, but also necessary to minimize false positives. To do this, we considered novel RE sites that would have been created if only one of the two requisite nt changes had occurred. As a comparison against the twin (AGCTGG to gGCcGG) substitutions used to create a novel HaeIII site (<u>GGCC</u>), we considered the frequency with which an adjacent HpaII site (CCGG) was formed that can also arise from a single nt substitution (AGCTGG to AGCcGG). Similarly, the frequency of a novel site arising from only one of the substitutions intrinsic to creating a BpmI site (CTGGAG via twin ATGGAG to ctGGAG substitutions) was calculated, though in this case an imaginary RE site was queried. Although the number of hypothetical false-positive WT reads increased from 19 to 144 in these single nt substitution scenarios (Table 2), the overall false-positive rate only increased from 0.06% to 0.4% (144 out of 33242 reads; Table 2), suggesting that an option for RE site creation via single nt changes could be a useful addition to future versions of TIARS Designer.

## DISCUSSION

A challenge of genome editing in the zebrafish model is the long feedback loop between the execution of a genome-editing perturbation and the identification of alleles carrying desired edits. For other model organisms with shorter life spans, such as *C. elegans* (3), it is sufficient to wait a few weeks (compared to 4 months for zebrafish) for the next generation and assess F_1_ progeny for germ line transmission. For mice, embryonic-stem cells allow for pre-screening of edits of interest. Our experience has been that the ability to assess somatic rates of desired edits in injected F_0_ embryos is critical for quality control and for efficiently comparing methods and variables. To this end, we routinely use the CRISPR-STAT fluorescent PCR-based approach for assessing the ability of gRNAs to produce standard indels (6).

We initially considered two methods for examining precise edits: subcloning of PCR amplicons and high-throughput sequencing. Our exploration of the former led to a waste of time due to a falsepositive artifact. We ultimately figured out that subcloning amplicons containing tandem repeats can produce altered sequences due to bacterial deletions or expansions of the repeat (15). Given that tandem repeats are a common outcome of Cas9/HDR procedures (19), we now avoid subcloning to evaluate Cas9/HDR outcomes and recommend that others use caution as well.

High-throughput sequencing is certainly a viable approach for assessing the somatic mutational spectrum among F_0_ embryos, and one that we use ourselves (manuscript in preparation). Here, we have presented our development of TIARS, an alternative approach that combines RFLP with CRISPR-STAT. While it is true that costly DNA analysis equipment is required for both TIARS and high-throughput sequencing, a growing number of companies perform these analyses for a fee and TIARS may be more time- or cost-efficient for small investigator-led projects than high-throughput sequencing. For instance, TIARS allows for a two-day turnaround from tissue harvest to data analysis, enabling researchers with limited husbandry space to quickly prioritize groups of animals of interest, such as F_0_ larvae from the same microinjected cohort as sampled specimens showing promising rates of somatic edits. Turnaround for high-throughput sequencing is longer, requiring at least one week, necessitating husbandry of a larger zebrafish population while awaiting results. Other pros and cons of TIARS vs. high-throughput-sequencing for detection of somatic edits have not been fully assessed, including the question of which method is more sensitive. The typical high-throughput sequencing error rate (below which no novel edits can reliably be detected) can reach 0.1% (20), whereas we have not yet determined the TIARS limits of detection.

Our group’s standard CRISPR/Cas9 practices were applied to this project and certain approaches to HDR editing were also adopted on faith. Among our standard practices is the use of commercially synthetized gRNAs. Standard laboratory synthesis of gRNAs requires a 5’GG pair for optimal yields (21). Certain workarounds, such as modification of gRNAs with 5’ NG or GN termini to 5’GG have been shown to be effective (22). Nonetheless, commercially synthesized gRNAs have no sequence constraints at all, allowing researchers to test any PAM-adjacent sequence of choice, such as the 5’TT gRNA we selected for the *atp7a^P1387S^* project.

Without having run any in-house quantitative comparisons against other ssODN designs, we have followed on faith the recommendation that ssODNs be complementary to the non-target strand and that the homology arms asymmetrically flank the predicted Cas9 cut site with a bias to the 5’ side (13). Also on faith, we have followed the principle of including PAM-site or gRNA-binding disruptions in our ssODNs, as theoretical inoculation against re-cutting of edited alleles by persisting gRNA/Cas9 (3). We did run a small-scale test of the proposal that phosophorothioate-end protection of ssODNs increases HDR editing efficiency as has been reported (16), with no supporting evidence in our hands, though we do not claim our single counter-example comprises a critical evaluation of the proposal.

Zebrafish *atp7a* is an essential gene (9), which rendered it impossible for us to apply yet another common recommendation for increasing HDR editing efficiency: that the gRNA cutting efficiency be maximized (3). The essential role of *atp7a* was indeed a constraint for us, even preventing it seemed, our successful use of a gRNA with a cutting efficiency as low as 24% on a scale that often exceeds 90%. The one gRNA that worked for us to establish germ-line transmission had the very low cutting efficiency of 7%. Our success using such an inefficient gRNA suggests that cutting efficiency can be sacrificed in certain contexts. The fact that this gRNA’s cut site was precisely at the location of the desired edit, another recommended HDR-editing design principle, seems likely to have contributed to our success (3). Of course genome-editing strategies that can alter coding sequence using gRNAs that target non-coding sequence and thus leave the protein unperturbed in most failed editing scenarios, such as the method of Hoshijima *et al.*, offer an alternative workaround for editing essential genes (5).

The exercise of manually searching for synonymous changes to create novel REs is both time consuming and error prone. Furthermore, the programming and implementation of TIARS Designer to solve this problem soon made us realize how far we had been from considering all ssODN design possibilities in any kind of unbiased or comprehensive manner. We subsequently have used this tool for a number of unpublished studies. Our foray into computational analysis of CRISPR/Cas9-based HDR editing also helped us more definitively establish that synonymous changes serve as excellent proxies for their non-synonymously changed neighbors, with extremely low false positive or false negative rates. These analyses further suggested that a lower bar of filtration allowing creation of novel RE sites via only one rather than two nt substitutions could be adequate. The question of whether supernumerary synonymous changes might in and of themselves affect HDR rates, is an outstanding question that we plan to experimentally address in future studies.

In closing, having conceived and developed TIARS and TIARS Designer as new tools in the genome-editing arsenal, we hope they will be useful to other researchers engaged in ssODN-based HDR gene editing projects and related approaches.

## DATA AVAILABILITY

TIARS <u>Designer, an open-source Python package available at</u> https://github.com/NICHD-BSPC/tiars-designer

## SUPPLEMENTARY DATA

Supplementary Table S1 and Figures S1-S4 have been provided in a separate file.

## ACKNOWLEDGEMENT

Thanks to Tianwei Li and Joseph R. Zoeller for assistance with high-throughput sequencing. Thanks to the aquatics staff of the NIH Central Aquatics facility and members of the Research Animal Management Branch for their excellent overall support and humanitarian care and oversight of the zebrafish used in this research. Thanks to Blake R. Carrington, Daniel A. Castranova, Andrew E. Davis, Tokunbor A. Lawal and Bradley S. Toms for technical support and intellectual input.

