## Supplemental Material for "Detection of precisely edited CRISPR/Cas9 alleles through co-introduced restriction-fragment length polymorphisms"

#### Supplemental Figure Legends

**Table S1. The six gRNAs and their separately determined cutting efficiencies considered for the *atp7a* editing projects.**

**Figure S1. Knock-in strategy for generating the *atp7a*$^{P1387S}$ allele.**
Schematic representation of the *atp7a* genomic structure (Intron 21, Exon 22 and intron 23, top line and bar). The ssODN donor template shown underneath is complementary to the nontarget strand containing the mutation (red arrow and pointed yellow star) adjacent to double strand break (lightning bolt) with synonymous changes introducing two RE sites - Hpy188I and HpyAV - resulting in 124 and 128 nt restriction fragments linked to primer F1, respectively. The WT amplicon has 331 nt linked to primer F1, which is reduced to 208 nt when digested with Hpy1881 due to an endogenous Hpy1881. PCR primers (black arrows).

**Figure S2. False positive "precise edit" detection when subcloning tandem repeats.** Sequencing outcomes from subclones derived from an $F_1$ heterozygote yielded a perfect edit ("mut"). This turned out to be a false positive artifact, which we discovered by direct sequencing (no subcloning) followed by peak disambiguation revealing a single tandem repeat at the target site of a portion of sequence from the ssODN that had been injected into the $F_0$ parent (same sequence as "mut + 48"). Note that not only is a perfect edit one of the sequence outcomes after subcloning ("mut"), but also a duplication of the tandem repeat (mut + 96). This phenomenon of deletion (mut + 48 > mut) and expansion (mut +48 > mut + 96) of direct repeats has been observed in bacteria and proposed to occur via slippage of the nascent strand, as shown in the schematic cartoon reproduced from: Bzymek, M. and Lovett, S.T. (2001) Instability of repetitive DNA sequences: the role of replication in multiple mechanisms. Proc Natl Acad Sci U S A, 98, 8319-8325.

**Figure S3. Further optimization of modified fluorescent PCR fragment analysis.** Optimization of fluorescent intensity (flu) was performed in parallel assays using the same fin biopsy lysate as Fig. 2 in combination with different variables. (A) Adding 5 μl vs.10 μl of fluorescent PCR product to RE-digests (or mock digests) of a final 15 μl volume from which 2 μl was subsequently sampled, mixed with HiDi formamide and denatured for capillary electrophoresis, using an injection time of 60s. 2.5 μl samples also functioned well (data not shown), but further increases in the proportion of RE-digested fluorescent PCR product in capillary electrophoresis samples led to failures (data not shown), presumably due to excess dilution of HiDi formamide and/or interference by excess salts, glycerol or protein from the RE digest. (B) Adding 10 μl PCR product to digests as in (A) and 23, 60 and 180s injection times. The combination of adding 10 μl PCR product to digests and 180s injection time yielded peaks with greater than 2-fold intensity over those shown in Fig. 2, where 5 μl and 60s parameters were used.

**Figure S4. Evidence of precise-edited *atp7aP*$^{1387S}$ mutant alleles in ssODN-injected 5 dpf $F_0$ embryos.**

**A. Right top panel**: control in the absence of RE. Indel fragments clustered around 331 nt, **Right middle panel**: Hpy1881 digestion of PCR product yields endogenous Indel fragment clustered around 208 nt and a proxy peak of precise editing at 124 nt. **Left panels**: re-scaled PCR/RFLP analysis. **Top**: control, **Middle**: Hpy188I and **Low**: HpyAV

**B.** Comparison of somatic editing efficiency between standard and phosphorothioate end-protected ssODN donor templates.

#### Table S1. Six gRNAs Considered for *atp7a* Editing Projects and their Cutting Efficiencies

| | **gRNA #1** | **gRNA #2** | **gRNA #3** |
|---|---|---|---|
| ***atp7a* T979I project** | **GGGAGGCCTGGAAGGCGAAG (AS strand)** | AGCGATGCAGAGGACAGTGA (AS strand) | GGTTGCGAGACCCAGAGAGC (AS strand) |
| **cutting efficiency** | **97%** | 7% | 30% |
| ***atp7a* P1387S project** | TTGGTTCTGCAGCCCTGGAT | GGCTGCAGAACCAACCCCAC | GGGGGTTGGTTCTGCAGCCC |
| **cutting efficiency** | 24% | 39% | 49% |

#### Fig. S1. Knock-in strategy for *atp7a*$^{P1387S}$ allele

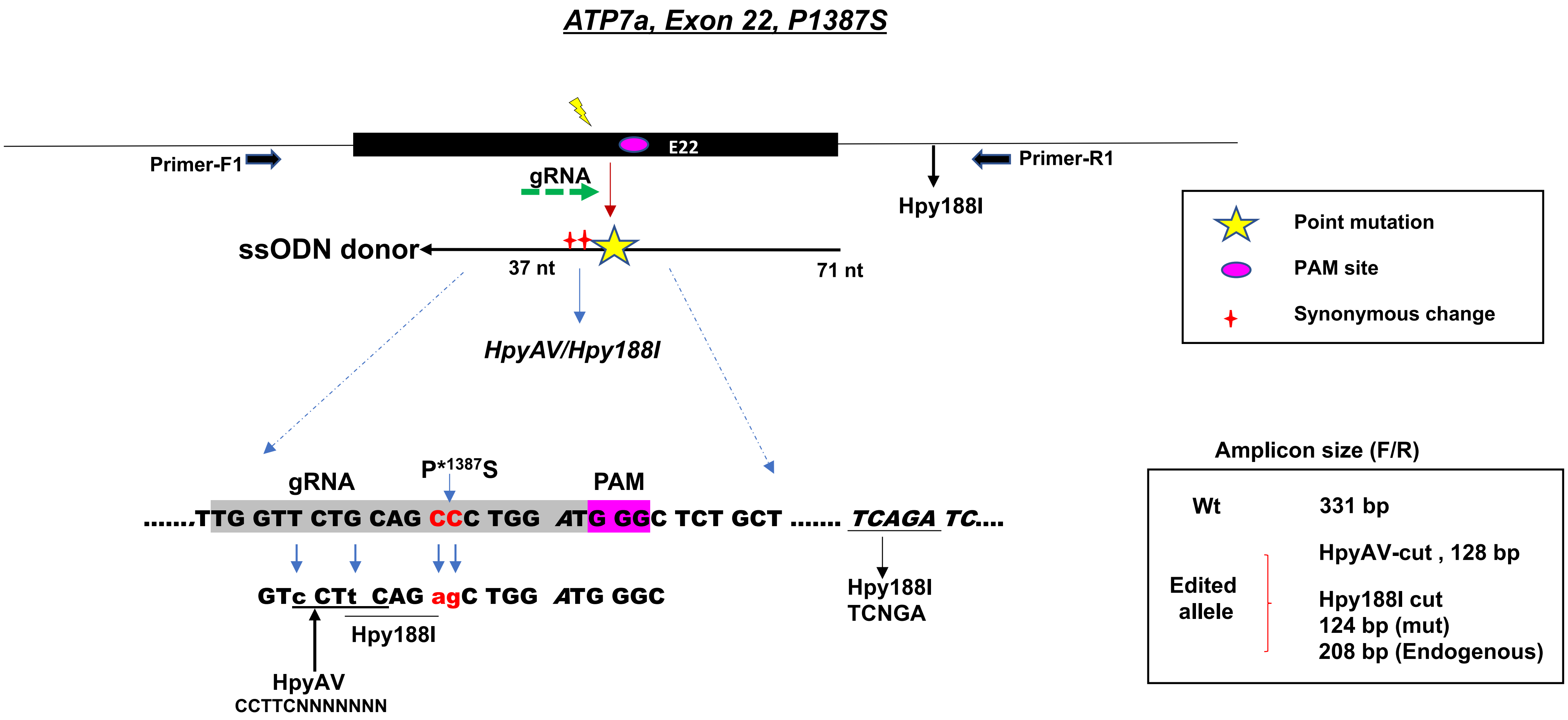

CATTTTAGCAGGAGGGAGGACAGCACTACAGACACAGAAGATAGAGCCATAGCAGCAGAGCCCATCCAGGACTGCAGGACCAACCCCACAGGC[illegible]

### Fig. S2. Subcloning DNA from $F_1$ heterozygote revealed that bacteria can create a false positive edit from a tandem repeat.

| | |
|---|---|
| **wt** | *AGCAGTGAATGGGGAAGATGCTACTTTAAAATATTCAGACTAAAGTGTTCT*<br>*GTTTTTTTTTTTTACTTGGTCAGGTTTCTATTTTGTCCTTGAACATATTGAATG*<br>*GAGAATGTGTTTCTCTGCAG* GGT TAT GAT AAG AGT ATC TCT GAA GCG<br>GAG GCT GTC ATC CGC TTC GCC TTC CAG GCC TCC ATC ACT GTC CTC<br>TGC ATC GCT TGC CCC TGC TCT CTG GGT CTC GCA ACC CCC ACA GCA<br>GTA ATG GTG GGC ACA GGG GTC GGA GCC CAG AAT GGA ATC CTC<br>ATC AAG GGA GAG CCA TTA GAG ATG GCA CAC AAG |
| **mut** | *AGCAGTGAATGGGGAAGATGCTACTTTAAAATATTCAGACTAAAGTGTTCT*<br>*GTTTTTTTTTTTTACTTGGTCAGGTTTCTATTTTGTCCTTGAACATATTGAATG*<br>*GAGAATGTGTTTCTCTGCAG* GGT TAT GAT AAG AGT ATC TCT GAA GCG<br>GAG GCT GTC ATC CGC TTC GCC TTC CAG GCa TCg ATa AtT GTC CTC<br>TGC ATC GCT TGC CCC TGC TCT CTG GGT CTC GCA ACC CCC ACA GCA<br>GTA ATG GTG GGC ACA GGG GTC GGA GCC CAG AAT GGA ATC CTC<br>ATC AAG GGA GAG CCA TTA GAG ATG GCA CAC AAG |
| **mut+48** | *AGCAGTGAATGGGGAAGATGCTACTTTAAAATATTCAGACTAAAGTGTTCT*<br>*GTTTTTTTTTTTTACTTGGTCAGGTTTCTATTTTGTCCTTGAACATATTGAATG*<br>*GAGAATGTGTTTCTCTGCAG* GGT TAT GAT AAG AGT ATC TCT GAA GCG<br>GAG GCT GTC ATC CGC TTC GCC TTC CAG GAT AAG AGT ATC TCT GAA<br>GCG GAG GCT GTC ATC CGC TTC GCC TTC CAG GCa TCg ATa AtT GTC<br>CTC TGC ATC GCT TGC CCC TGC TCT CTG GGT CTC GCA ACC CCC ACA<br>GCA GTA ATG GTG GGC ACA GGG GTC GGA GCC CAG AAT GGA ATC<br>CTC ATC AAG GGA GAG CCA TTA GAG ATG GCA CAC AAG |
| **Mut+94** | *AGCAGTGAATGGGGAAGATGCTACTTTAAAATATTCAGACTAAAGTGTTCT*<br>*GTTTTTTTTTTTTACTTGGTCAGGTTTCTATTTTGTCCTTGAACATATTGAATG*<br>*GAGAATGTGTTTCTCTGCAG* GGT TAT GAT AAG AGT ATC TCT GAA GCG<br>GAG GCT GTC ATC CGC TTC GCC TTC CAG AAG AGT ATC TCT GAA GCG<br>GAG GCT GTC ATC CGC TTC GCC TTC CAG GAT AAG AGT ATC TCT GAA<br>GCG GAG GCT GTC ATC CGC TTC GCC TTC CAG GCa TCg ATa AtT GTC<br>CTC TGC ATC GCT TGC CCC TGC TCT CTG GGT CTC GCA ACC CCC ACA<br>GCA GTA ATG GTG GGC ACA GGG GTC GGA GCC CAG AAT GGA ATC<br>CTC ATC AAG GGA GAG CCA TTA GAG ATG GCA CAC AAG |
| **Mut+38** | *AGCAGTGAATGG*GGAAGATGCTACTTTAAAATATTCAGACTAAAGTGTTCT<br>GTTTTTTTTTTTTACTTGGTCAGGTTTCTATTTTGTCCTTGAACATATTGAATG<br>GAGAATGTGTTTCT CTGCAG GGT TAT GAT AAG AGT ATC TCT GAA GCG<br>GAG GCT GTC ATC CGC NNC CCT CCG GTG GNG TTC GCT GNG CGN<br>NGG CTG TCT CCG CTT CC TTC CAG GCa TCg ATa AtT GTC CTc TGC ATc<br>GCT TGC CCC TGC TCT CTG GGT CTC GCA ACC CCC ACA GCA GTA ATG<br>GTG GGC ACA GGG GTC GGA GCC CAG AAT GGA ATC CTC ATC AAG<br>GGA GAG CCA TTA GAG ATG GCA CAC AAG |

#### Slippage model for genetic rearrangements

A slipped alignment of the nascent strain with respect to template can lead to the deletion or expansion of a directly repeated sequence and any intervening sequence

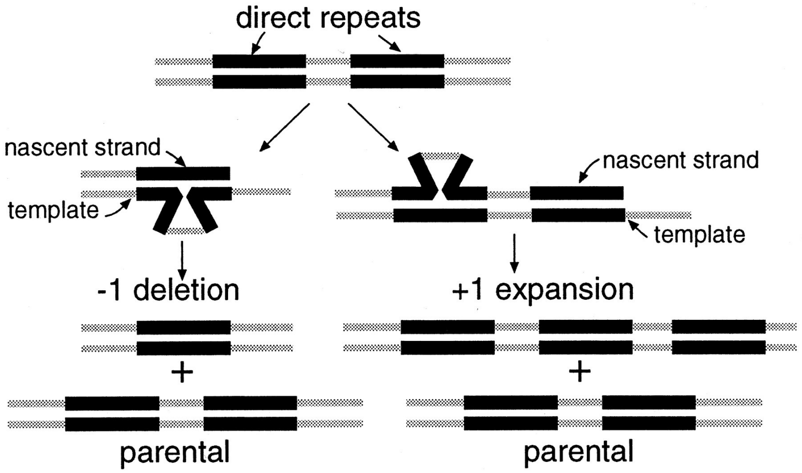

Fig. 1 of Bzymek, M. and Lovett, S.T. (2001) Instability of repetitive DNA sequences: the role of replication in multiple mechanisms. Proc Natl Acad Sci U S A, 98, 8319-8325. Copyright (2001) National Academy of Sciences, U.S.A.

**Beware of recombining tandem repeats during subcloning**

Fig. S3. Optimization of modified fluorescence PCR /RFLP fragment analysis using *atp7a*$^{T979I}$ as the template

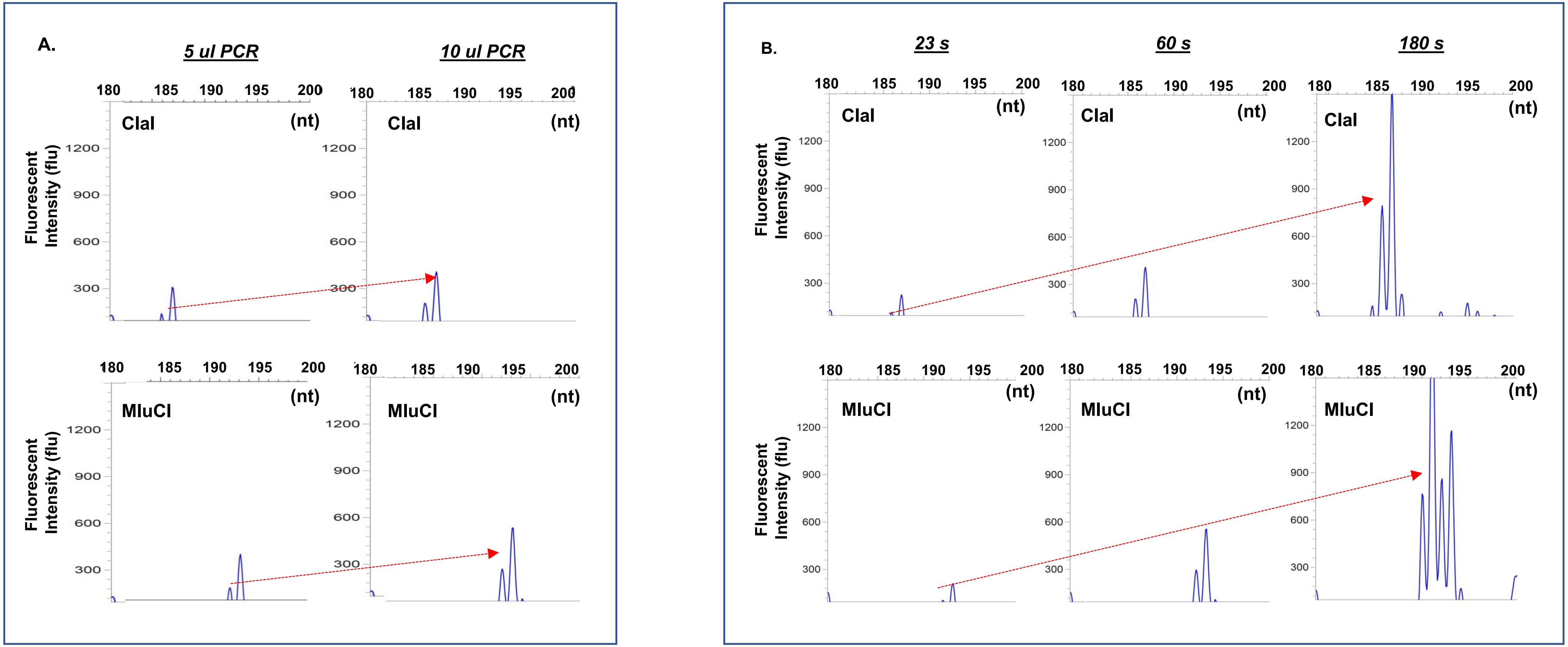

#### Fig. S4. Identification of precise-edited *atp7a*$^{P1387S}$ alleles (5 dpf $F_0$ embryos) by PCR/RFLP

A.

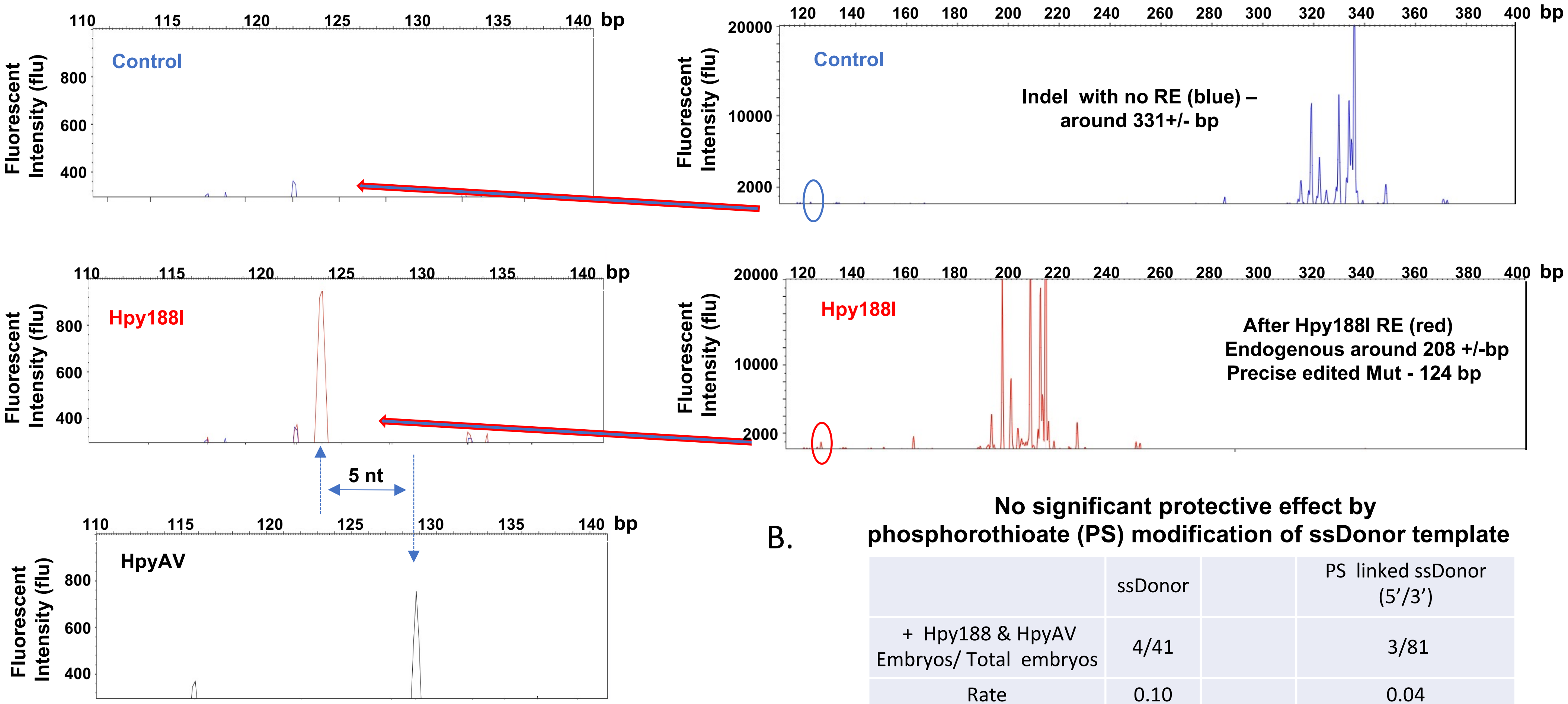

B.

**No significant protective effect by phosphorothioate (PS) modification of ssDonor template**

| | ssDonor | | PS linked ssDonor (5'/3') |
|---|---|---|---|
| + Hpy188 & HpyAV Embryos/ Total embryos | 4/41 | | 3/81 |
| Rate | 0.10 | | 0.04 |
